# Bacterial envelope damage inflicted by bioinspired nanospikes grown in a hydrogel

**DOI:** 10.1101/2020.03.28.013797

**Authors:** Sandra L. Arias, Joshua Devorkin, Jessica C. Spear, Ana Civantos, Jean Paul Allain

## Abstract

Device-associated infections are one of the deadliest complications accompanying the use of biomaterials, and despite recent advances in the development of anti-biofouling strategies, biomaterials that exhibit both functional tissue restoration and antimicrobial activity have been challenging to achieve. Here, we report the fabrication of bio-inspired bactericidal nanospikes in bacterial cellulose and investigate the mechanism underlying this phenomenon. We demonstrate these structures affects preferentially stiff membranes like those in Gram-positive bacteria, but exhibit cytocompatibility towards mammalian cells, a requisite for tissue restoration. We also reveal the bactericidal activity of the nanospikes is due to a pressure-induced mechanism, which depends on the cell’s adherence time, nanospike’s geometry and spacing, cell shape, and mechanical properties of the cell wall. Our findings provide a better understanding of the mechanobiology of bacterial cells at the interface with nanoscale structures, which is fundamental for the rational design bactericidal topographies.

## 1. Introduction

A characteristic hallmark in the pathogenesis of device-associated infections is the formation of biofilms at the biomaterial surface^1^. Biofilms are formed by bacterial communities enshrouded by exopolymers and extracellular DNA and are critical to the virulence of bacterial and fungal pathogens. Compared to a planktonic lifestyle, bacteria in biofilms are highly resistant to clearance by host immune responses and are recalcitrant to antibiotic treatment^1,2^. Once established at the biomaterial surface, bacteria can disseminate to other body sites and elicit severe secondary infections that contribute dramatically to patient morbidities and life-threatening events^3,4^. Despite the efforts in the field, infections caused by biofilms continue to be one of the leading causes of nosocomial infections, with an economic impact comparable to that of major diseases such as breast and colorectal cancer^5^. On top of this, current estimates project their incidence will increase significantly by 2050 as a result of antibiotic resistance^2,5^.

The first step in biofilm growth is known to be dictated by the physiochemical properties of the surface. It occurs as a result of the surface colonization by opportunistic pathogens that constitute the healthy microflora in mucosal membranes and skin or are present on the environment^6,7^. Since bacterial adhesion is critical to biofilm formation, multiple therapeutic strategies have focused on eliminating the undesirable presence of bacteria on biomaterial surfaces by different means, including the diffusive release of antibacterial agents and the use of anti-biofouling coatings^3^. However, those strategies face several drawbacks such as the lack of selectivity, suboptimal doses at the infected site, potential toxic accumulation at organs like kidney and spleen, and in the case of coatings, delamination and lack of durability upon mechanical stress^8,9^. Currently, the continuous replacement or removal of the biomaterial is still the most effective way to eradicate bacterial biofilms; however, this entails massive trauma, especially for a permanent internal prosthesis^1,3^.

Recent studies have demonstrated that interfaces with micro and nanoscale features exhibit increased biocompatibility towards mammalian cells while creating a hostile niche for bacterial colonization and growth. For example, micro and nanotopographical cues have been shown to encourage cellular growth and spreading, raising the likelihood of biomaterial integration with the host tissue^10,11^. Contrary to the positive effect on eukaryotic mechanotransduction, analogous topographical cues can limit bacterial adhesion and be detrimental to bacteria and their biofilms^8^. For example, the wing surface of insects like dragonfly and cicada species and the skin in gecko lizards are decorated with nanoscale protrusions able to mechanically disrupt bacterial cells within minutes^12–15^. The bactericidal activity of those naturally occurring nanotopographies has been shown to be limited to cells that made direct contact with the topography and to be independent on the surface chemistry^14,16^.

Compared to chemical-based bactericidal agents, bactericidal nanotopographies (e.g., nanopillars, nanospikes, and nanocones) are not consumed metabolically, are not leached to surrounding tissues, and do not contribute to bacterial resistance^8,17^. However, the mechanism by which nanoscale protrusions achieves its bactericidal activity remains poorly understood. Various processes mediating the action of those bactericidal topographies have been proposed, and those include the mechanical stretching of the bacterial envelope in contact with the topography and a shear-stress mechanism due to cell motility^13,18,19^. In the first scenario, the loss of bacterial membrane integrity is reasoned to occur in the region that suspends between adjacent nanoscale protrusions^18^. This model also proposes that soft membranes like those in Gram-negative bacteria are more predispose to rupture because they can easily absorb and conform to the topography compared to the thicker walls in Gram-positive species, which are thought to be more resistant to mechanical disruption^18^. Under this premise, one may expect the bactericidal activity to increase as the thickness of the nanocone apex increases. However, experimental evidence has demonstrated an opposite trend, in which the bactericidal activity increases as the nanoscale protrusions become sharper and taller^16^. Similarly, regarding a shear-stress mechanism linked to cell motility, experiments have shown that bactericidal nanotopograhies can also damage the membrane in microorganisms that lack any propulsion machinery^12,20^. Furthermore, and despite their promising functionality, up to now naturally occurring bactericidal topographies like those in the dragonfly and cicada wings have been mimicked on materials like poly(methylmethacrylate)^21^, titanium^19^, diamond^22^, and silicon^16^. However, much less work has been conducted on providing those attributes to clinically relevant hydrogels and very compliant polymers. Specifically, hydrogel nanopatterning has been limited by the complexity of obtaining patterns with precise control of the size that can also resist aqueous immersion^23^.

Here, we report the fabrication of bio-inspired bactericidal nanospikes fabricated in bacterial cellulose (BC). BC is a sustainable, biodegradable, and biocompatible hydrogel synthesized by microorganisms of the genus *Komagataeibacter*, which consists of glucan chains forming a porous structure^24^. BC is commonly employed as a candidate for cell scaffolds, artificial skin, vascular substitutes, and wound dressing^25–28^. Singly charged argon ions were used to induced pattern formation in two sets of BCs with differences in ribbon width, resulting in the evolution of high-aspect-ratio (HAR) nanostructures with “nanospike” appearance, but different feature size and spacing. In marked contrast with previous reports on bactericidal nanotopographies, we found that the nanospikes affected preferably stiff membranes like those in Gram-positive bacteria; however, this attribute also depended on the cell’s adherence time, nanospike’s geometry and spacing, cell shape, and mechanical properties of the cell wall. Atomic force microscopy (AFM) further confirmed the higher susceptibility of *B. subtilis* towards mechanical disruption. Moreover, the loading forces obtained by AFM were in quantitative agreement with computational simulations of a pressure-induced bactericidal mechanism.

## 2. Results

We induced nanopattern formation in bacterial cellulose (BC) by treating the hydrogel with argon (Ar^+^) ions at an energy of 1 keV, ion doses of 1×10^18^ cm ^-2^, and at a normal angle of incidence. Nanoscale features with “nanospike” in appearance were obtained by using two sets of bacterial cellulose, that is, a commercially available bacterial cellulose (CoBC) and a lab-grown bacterial cellulose (LgBC), which differ from each other in ribbon widths, i.e., 65±12 over 22±9 nm for the CoBC and LgBC, respectively (**Figure 1a**). These differences in ribbon width yielded nanospikes with variable size, but approximately the same aspect ratio (height-to-width) of ≈ 5. The commercially available argon-treated bacterial cellulose (CoArBC) developed tall nanospikes with an average height of 414 ± 82 nm and width at the nanocone’s bottom of 73 ± 16 nm. In contrast, the lab-grown argon-treated bacterial cellulose (LgArBC) grew shorter nanospikes with an average elevation of 222 ± 65 nm and a width of 43 ± 10 nm. The size at the nanocone apex was 50 ± 13 nm and 36 ± 8 nm in diameter for the CoArBC and LgArBC, respectively. We have recently revealed the mechanism of ion-induced nanopatterning in BC^29^. By studying the transformation of the material as a function of the ion dose, we discovered the onset of nanopattern formation was linked to the growth of carbon-rich clusters that were graphite-like in composition^29^. Pattern evolution was accompanied by changes in the Young modulus of both celluloses, which increased from 6.1 to 8.8 MPa in CoArBC, and from 1.5 to 9.8 MPa in LgArBC **(Figure 1c)**. However, there were no statistically significant differences in Young’s modulus between CoArBC and LgArBC, indicating that both celluloses developed a similar crosslinking density upon Ar+ treatment. Furthermore, CoArBC and LgArBC preserved their hydrophilic nature despite the increase in surface roughness, due to the presence of hydroxyl and carboxyl groups at the material surface^29^.

**Figure 1.**
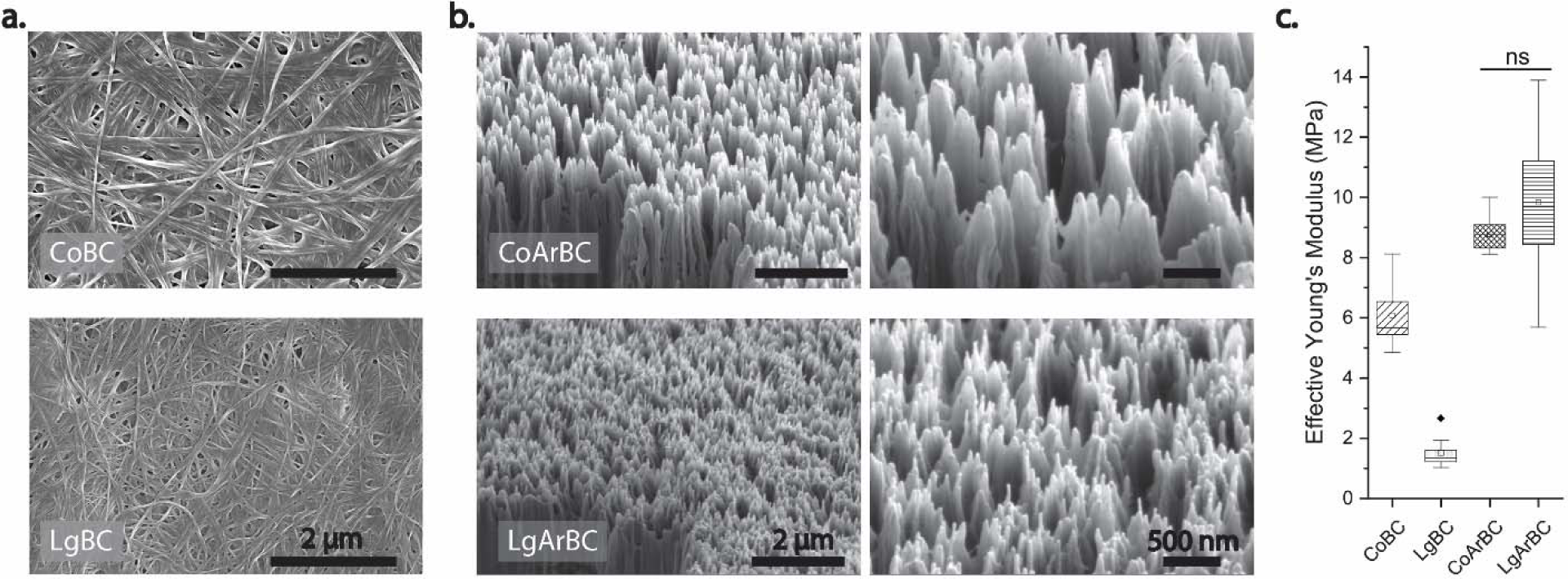
Nanospike size and spacing depend on the cellulose native ribbon width. Scanning electron micrographs (SEM) depicting the interlaced ribbon structure of pristine commercially available BC (CoBC) and lab-grown BC (LgBC) before **(a)** and after **(b)** treatment with singly charged argon (Ar^+^) ions. The right panels in **(b)** are high-resolution scans of the argon-treated BC. **(c)** Effective Young’s modulus for pristine and nanostructured cellulose, measured in ultrapure water with a soft material indenter. In all box plots, lines represent the median, small squares indicate the mean, dots are outliners, whiskers show the standard deviation (SD). No significant (ns) differences between groups.

Bioinspired by the dragonfly and cicada wings, we next investigated the bactericidal activity of the nanospikes against *Escherichia coli, Bacillus subtilis*, and *Staphylococcus aureus*. We examined bacterial viability and biofilm formation after 1 and 24 h of incubation with the experimental samples using a Live/Dead *BacLight™* bacterial viability kit and crystal violet staining. As a control, we used pristine CoBC. Short incubation times (i.e., 1 h) were used to get a better estimation of the cell viability since the propidium iodide in the *BacLight™* has also been shown to stain the extracellular nucleic acids (e.g., extracellular DNA) commonly found in mature biofilms^30^. Moreover, the bactericidal activity of nanoscale topographies has been reported to occur at a time scale of minutes^14^. Importantly, cofounding factors in envelope disruption stemming from capillary forces were minimized by performing all our analysis in an aqueous environment (minimal media or phosphate-buffered saline (PBS) solution). Among the bacterial species used, *E. coli* and *S. aureus* have been associated with a wide range of human diseases, including urinary tract infections, bacteremia, cystic fibrosis, and liver abscesses and have become increasingly resistant to antibiotic treatment^31,32^. *B. subtilis* was selected as a model microorganism to correlate the mechanical properties of the bacterial envelope, e.g., the peptidoglycan thickness, to the extent of cell damage inflicted by the nanotopography. The cell envelope, or the outer surface in bacteria, is a multilayer structure that varies in composition and structural organization among Gram-negative and Gram-positive species^33^. The envelope in Gram-negative bacteria (e.g., *E. coli*) consists of a cytoplasmic and outer membrane separated by a thin peptidoglycan wall of about 2.5-6.5 nm thick. Conversely, the envelope in Gram-positive bacteria (e.g., *B. subtilis* and *S. aureus*) compromises a single cytoplasmic membrane with a thick peptidoglycan wall of 19-33 nm. The peptidoglycan is the primary structural polymer of the bacterial cell wall and is thought to make the Gram-positive species significantly stiffer than their Gram-negative counterparts^34^. Based on its structural composition, thick peptidoglycan walls have also been suggested to confer higher resistance to Gram-positive species against disruption by nanoscale protrusions^16^.

We found that the viability of *B. subtilis* decreased significantly on both CoArBC and LgArBC when compared to the control samples at 1 h incubation (**Figure 2a**). Unexpectedly, the percentage of viable *E. coli* was significantly higher than *B. subtilis* in both CoArBC and LgArBC. We also found that the nanospikes in CoArBC yielded a more significant percentage of dead *B. subtilis* cells as opposed to the denser nanospike array in LgArBC (**Figure 2b**). However, the nanospike’s size and spacing had little effect on the *S. aureus* viability, which was not significantly different between control and nanostructured celluloses (**Figure 2a, b**). These results suggest that Gram-negative cells may be more resistant to mechanical disruption than their Gram-positive counterparts. Yet, among Gram-positive species, rod-shaped bacteria (e.g., *subtilis*) are more significantly affected than spherical-shaped ones (e.g., *S. aureus*).

**Figure 2.**
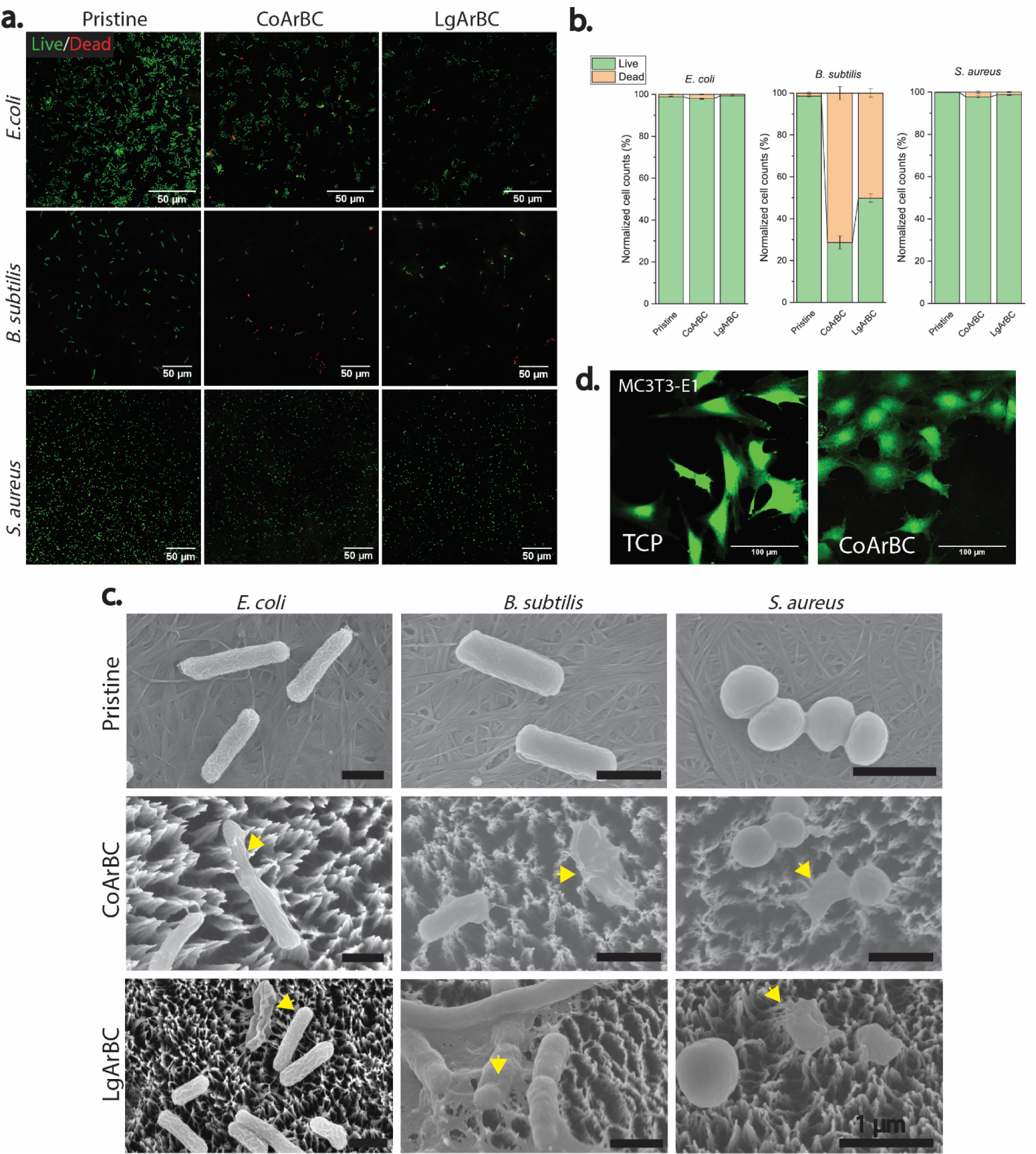
The nanostructured celluloses disrupted the envelope preferentially in *B. subtilis* but did not compromise the integrity of murine pre-osteoblasts. **(a**) Confocal laser scanning microscopy images of live (green) and dead (red) *E. coli, B. Subtilis*, and *S. aureus* stained with a live/dead *Bac*Light™ viability kit and cultured for 1 hour on CoArBC and LgArBC; **(b)** percentages of the dead to living bacteria normalized to the total cell counts per unit area. The bar plots and error bars show the mean ± SEM, respectively. **(c)** Scanning electron micrographs show the typical shapes of viable bacteria on the pristine BC and lysed bacterial cells (yellow head arrows) on CoArBC and LgArBC substrates. **(d)** MC3T3-E1 cells stained with a Live/Dead Viability/Cytotoxicity Kit seeded on tissue-culture polystyrene (TCP, control sample) and CoArBC. The green fluorescent staining demonstrates the plasma membrane integrity on those cells on both substrates.

Scanning electron microscopy was used to visually inspect the morphology of the different bacterial species in contact with the experimental samples (**Figure 2c**). In the pristine BC, the cells displayed the typical shape of a functional bacterium, while several of those cultured on the nanostructured celluloses appeared flat, indicating that the cellular contents leaked out (**Figure 2c and Supplementary Figure S1a**). Those observations are consistent with reports made by other authors using either naturally occurring bactericidal topographies or replicas derived from the natural counterparts^12,13,16,20^. Moreover, compared to *E. coli* and *B. subtilis* that laid in direct contact with the nanospike apexes, *S. aureus* could colonize the gap separating adjacent structures, particularly in the CoArBC (**Supplementary Figure S1b**). Most importantly, the nanospikes did not compromise the viability of murine pre-osteoblasts as determined via a Live/Dead Viability/Cytotoxicity staining. In the CoArBC, the morphology of MC3T3-E1 cells was similar to that in tissue culture polystyrene (TCP), which was used as a control surface (**Figure 2d**). Furthermore, in both CoArBC and TCP, the MC3T3-E1 demonstrated intracellular esterase activity and plasma membrane integrity, with near 100% viability, implying that the nanostructured BC is cytologically compatible.

Biofilm assessment using crystal violet staining after 24 h of incubation revealed a significant increase in biofilm mass accumulation on CoArBC and LgArBC compared to the pristine BC for both *E. coli* and *B. subtilis*, except in *S. aureus*, where the biofilm mass remained similar to the control (**Figure 3a**). A closer examination using the live/dead *Bac*Light™ viability kit showed that the accumulated biofilm mass on the nanostructured celluloses was mostly made up of dead cells as depicted by the intense red fluorescence signal coming from the propidium iodide nucleic acid stain. This red staining was not observed in the controls for any of the strains tested, meaning this staining corresponded to nucleic acids inside the bacteria with compromised membranes and not from extracellular DNA (**Figure 3b**). In sharp contrast with the observations made at 1 h incubation, *E. coli* biofilms on CoArBC, and to a lower extent in the LgArBC, also showed a significant accumulation of dead bacteria compared to the unmodified BC (30% vs. 0.5% of dead cells) (**Figure 3c**). Similarly, the percentage of dead *B. subtilis* cells increased by about 10% at 24h for both CoArBC and LgArBC compared to 1 h incubation, but S. *aureus* biofilms remained viable and not significantly different from the control. Consistent with shorter incubation times, CoArBC showed a more significant bactericidal activity than LgArBC. Moreover, the bactericidal activity of the nanospikes was independent of the surface chemistry, since a 30 nm gold coating did not abolish this attribute in CoArBC when tested against *B. subtilis* (**Supplementary Figure S2**). In agreement with other reports, this result suggests that the contact killing mechanism of nanoscale protrusions is independent of the material’s chemistry^16,20^. Overall, these observations confirmed a more significant susceptibility of *B. subtilis* towards the nanospikes in the BC, which increased over time even for *E. coli* and was most conspicuous in CoArBC than in the tightly packed nanospikes present on LgArBC.

**Figure 3.**
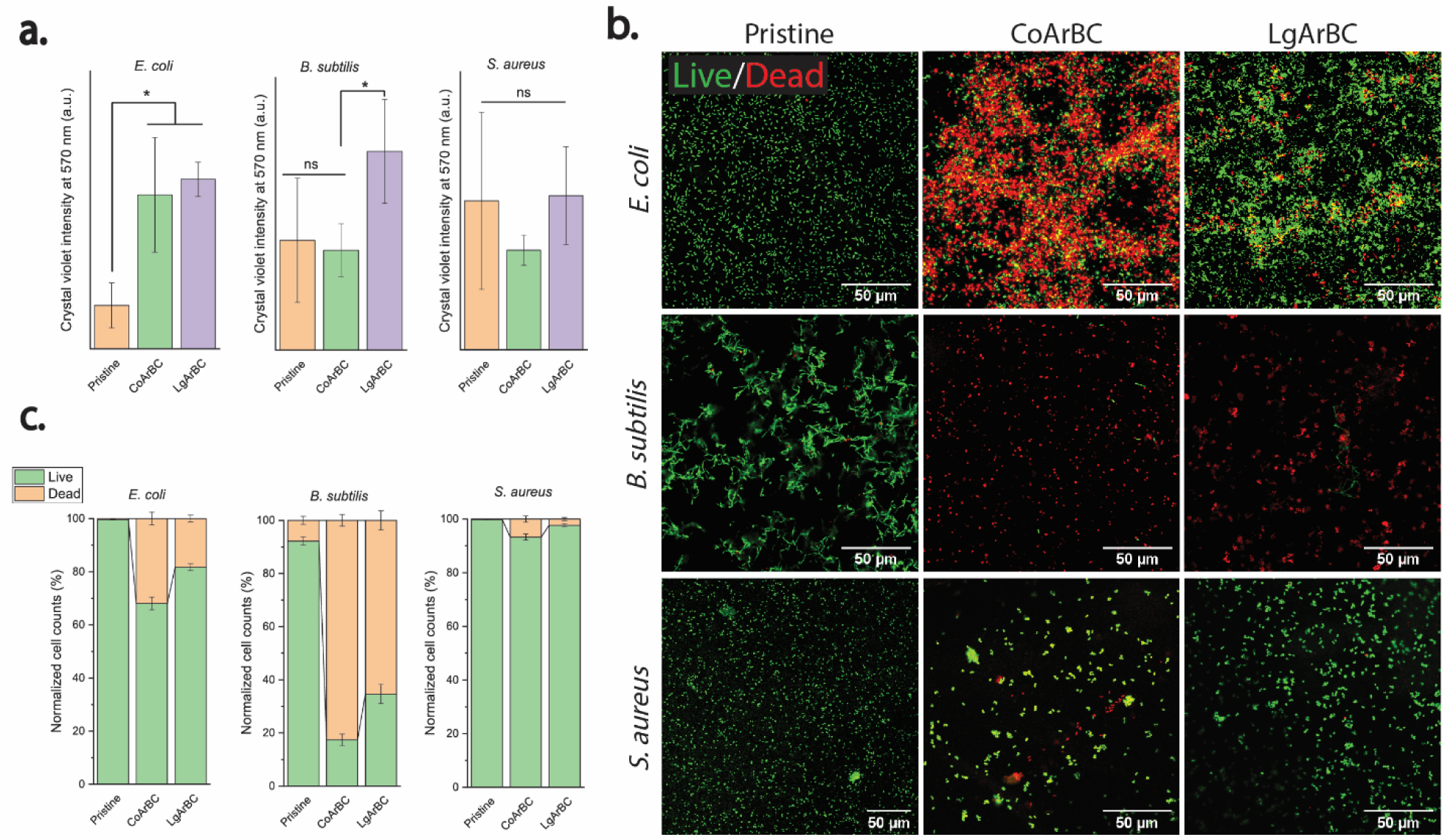
The bactericidal activity of the nanostructured celluloses depends on the cell’s adherence time. **(a)** Quantification of *E. coli, B. subtilis*, and *S. aureus* biofilms using Crystal Violet after 24 h incubation with the experimental samples. The asterisk denotes significant differences (*p*<0.05). Bar plots correspond to the mean ± SD. **(c)** Confocal laser scanning micrographs of *E. coli, B. subtilis*, and *S. aureus* biofilms stained with *BacLight* Live/Dead kit. Images for *E. coli* and *B. subtilis* correspond to 3-D projections. **(d)** Quantification of the dead to living cells obtained from (b). The bar plots show the percentage of live and dead cells per unit area ± SEM.

A cross-sectional examination of *E. coli* and *B. subtilis* at the interface with the nanostructured celluloses revealed that the nanospikes indented those strains in the region immediately above the nanocone apex (yellow arrowheads in **Figure 4**), whereas changes in the thickness of the bacterial envelope suspending between adjacent nanospikes were not visible. The direct contact between the apex of HAR nanostructures with biological membranes has already been implicated in the penetration of mammalian cells by silicon nanowires^35,36^ and has also been observed in yeast cells in contact with dragonfly wings^12^. This suggests the nanospikes may pre-deform the bacterial envelope before its mechanical disruption. Remarkably, the indentation of the bacterial envelope was several nanometers deep in both *E. coli* and *B. subtilis* and approximately equal to 160±66 nm and 157±5 nm, respectively, in CoArBC.

**Figure 4.**
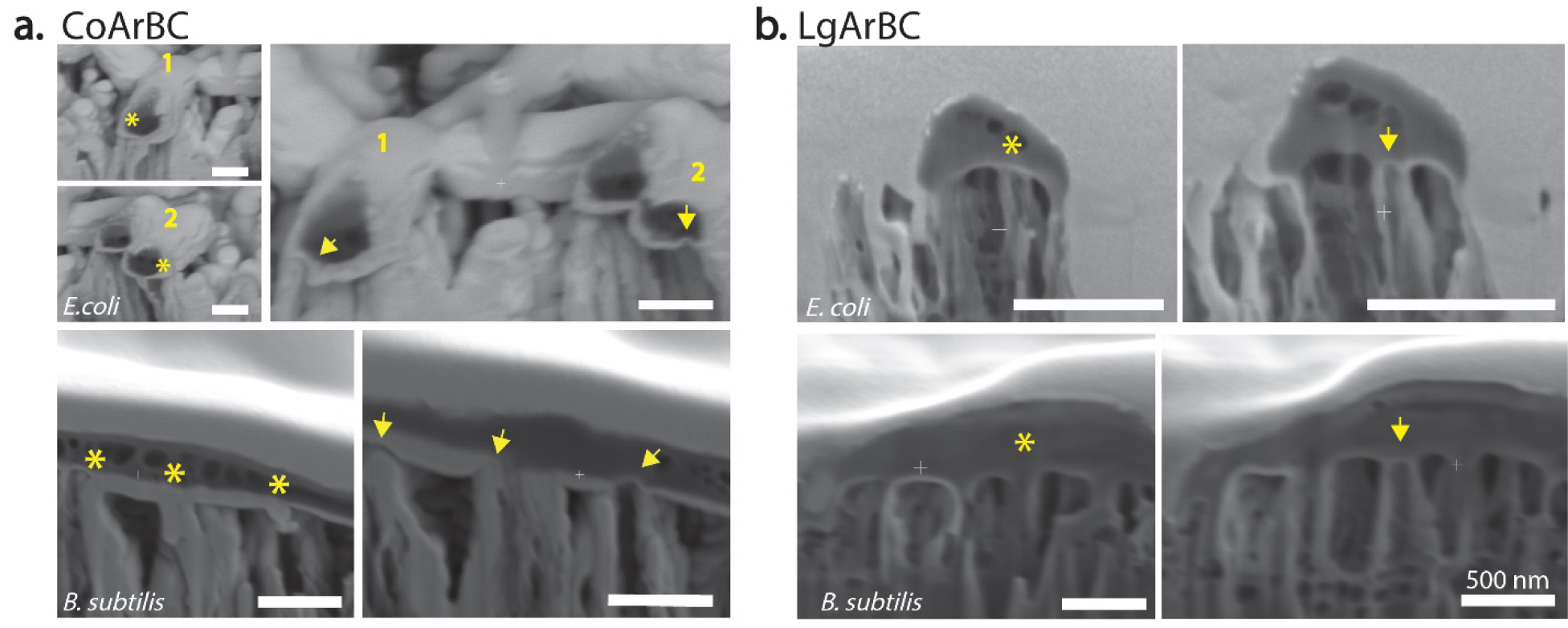
The nanospike apexes indented *E. coli* and B. *subtilis* by several nanometers deep. Cross-sections of *E. coli* and *B. subtilis* in contact with **(a)** CoArBC and **(b)** LgArBC. The asterisks and arrowheads denote before and after the cross-section, respectively. Notice the indentation at the cell wall left by the nanospike (indicated by the yellow arrows).

To understand the preferential loss of integrity in *B. subtilis* compared to *E. coli*, we measured the indentation force experienced by those cells at the interface with the nanospikes. Specifically, we looked for quantifying the loading force required to breakthrough *E. coli* and *B. subtilis* with nanoprobes of about 19±3 and 30±9 nm radii, which are comparable to the apex in the nanostructured celluloses (**Figure 1a**). **Figure 5a** shows representative force volume maps obtained for each bacterium using the 19 nm tip radius nanoprobe, whereas **Figure 5b** depicts two representative force curves with the typical signatures of breakthrough events of the bacterial envelope extracted from those force volume maps (green dots in **Figure 5a**). Notice that as the cantilever advances toward the surface (extension curve, **Figure 5b**), the lipid bilayer is elastically deformed up to a threshold value where the membrane cannot further withstand applied force. This loading force (or breakthrough force) is characterized by a discontinuity or “a jump” as the cantilever advances through the cell thickness up to reach the substrate^37^. In contrast, discontinuities during the retraction of the cantilever (retraction curve, **Figure 5b**) are linked to adhesion events between the cell and the cantilever tip.

**Figure 5.**
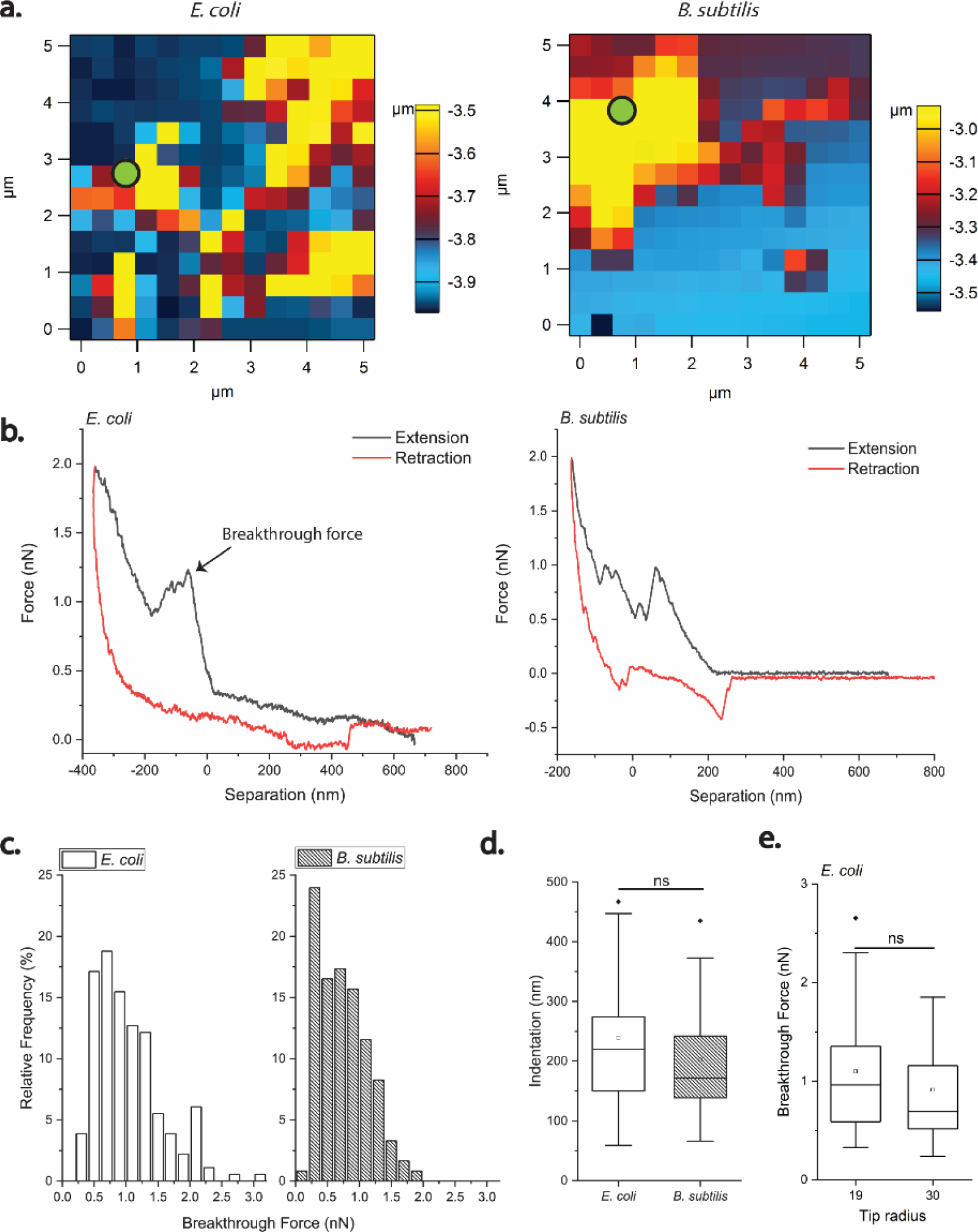
*B. subtilis* is penetrated by AFM nanoprobes at lower loading forces than *E. coli*. **(a)** Force-volume images of *E. coli* and *B. subtilis* obtained with a cantilever tip of 19±3 nm in radius. Each image contains 14×14 pixels, and each pixel corresponds to a force curve. The blue pixels denote the substrate (silicon), whereas the yellow and red pixels correspond to living bacterial cells. The force volume map was obtained on a random area of 5×5 µm^2^. **(b)** Representative force curves for *E. coli* and *B. subtilis*, whose position in the force map is indicated by green dots. The first discontinuity or “jump” in the extension curve corresponds to the breakthrough force. The baseline in both the extension/retraction curves was corrected using FRAME (see materials and methods) before determining the breakthrough force. **(c)** Relative frequency histograms showing the distribution of the breakthrough force in *E. coli* and *B. subtilis* for loading forces of up to 5 nN. Two-sample Kolmogorov-Smirnov test indicated both data sets came from different distributions (*p-value* < 8.9 x10^−5^). **(d)** Before rupture, the bacterial envelope in both *E. coli* and *B. subtilis* was deformed by several nanometers; however, this value was not significantly different between the two cell types. **(e)** The breakthrough force does not change significantly by indenting *E. coli* with a thicker cantilever tip. Not significant: ns.

For our analysis, we selected the first breakthrough event, as it has been demonstrated to represent the penetration of the first lipid bilayer as the cantilever moves through the thickness of living mammalian cells and multi-stacked membranes^38,39^. To determine the loading force required to penetrate the envelope in *E. coli* and *B. subtilis*, we systematically obtained more than 50 force volume maps of 196 pixels each, yielding more than 9800 force curves per cell type. We then selected the curves that unambiguously displayed the breakthrough signatures previously characterized by other authors on synthetic lipid stacks and living cells^38,40^. By using the nanoprobe of ≈ 19 nm tip radius, we found that loading forces below 3 nN were enough to breach the wall in both *E. coli* and *B. subtilis* (**Figure 5b**). This breakthrough force is comparable to the reported for *Salmonella typhimurium* indented by nanoprobes with tip radii ranging between 25 and 35 nm^40^. Importantly, the wall in *B. subtilis* failed in average at lower loading forces (0.74 ± 0.38 nN) than in *E. coli* (1.04 ± 0.52 nN), and this difference was significant. **Figure 5c** shows the relative frequency histograms depicting the breakthrough force distributions for both *E. coli* and *B. subtilis*. Two-sample Kolmogorov-Smirnov test applied on these data sets further confirmed they came from different distributions (*p-value* < 8.9 ×10^−5^). These results correspond well to the higher susceptibility of *B. subtilis* to the nanospikes in the argon-treated BC we quantified by live/dead fluorescence staining at 1 and 24 h. Compared to *B. subtilis*, however, the probability to breakthrough *E. coli* at loading forces in the range of 2 to 5 nN displayed a higher variability and wider distribution. To test the possibility of a spatially dependent breakthrough force in *E. coli*, we indented a single *E. coli* cell along its longitudinal axis (**Supplementary Figure S3**). We found that for a loading force of 2nN, the region close to the cell poles deformed elastically, whereas the cell failed preferentially at the center (**Supplementary Figure S3**). We also detected the indentation depth was not significantly different between *E. coli* and *B. subtilis*, which was 239±124 nm and 202±93 nm, respectively (**Figure 5d**). Moreover, this indentation depth was comparable to the deformation of the cell envelope detected at the interface with the nanospikes in CoArBC (160±66 nm in *E. coli* and 157±5 nm in *B. subtilis*) (see **Figure 4**).

Interestingly, breakthrough force in *E. coli* was not significantly different when the cell was indented by a thicker cantilever tip (30 ± 9 nm) (**Figure 5e**), which is consistent with observations previously made by other authors^38,40^. Also, we did not observe an apparent decrease in height when *E. coli* or *B. subtilis* were penetrated repeatedly, indicating that upon cantilever retraction, the bacterial envelope was able to recover its integrity. In opposition to this observation, the indentation of fixed *E. coli* produced large deformations in the cell, which caused the bacterium to shrink against the substrate (**Supplementary Figure S4**). This behavior agrees with observations made in fixed *S. typhimurium*^40^, indicating that the mechanical properties of dead bacteria differ substantially from those in living cells.

## 3. Discussion

Nanoscale protrusions like those found in the wing surface of cicada and dragonflies have been proven to be antimicrobial against bacteria and yeast cells, but a mechanistic understanding of the underlying phenomenon is still debatable. Moreover, previous work has focused on mimicking naturally occurring bactericidal topographies on acrylic resins, metals, and semiconductors, but little progress has been made on providing those attributes to clinically relevant hydrogels and very compliant polymers. In this study, we were able to confer bactericidal activity to a BC by altering its topography at the nanoscale. Nanopatterning of the BC was achieved by treating the cellulose with low energy Ar^+^ ions, which resulted in the formation of HAR nanostructures with “nanospike” in appearance. Moreover, we obtained nanospikes with distinct size but comparable aspect ratio by using two sets of BC pellicles with different ribbon width, which were designated as CoArBC and LgArBC, based on the source of the pristine hydrogel. The ion-induced nanopatterning of BC differs from the pyrolysis of cellulose in that the polymer transformation is limited to the material surface and is driven by an athermal collisional radiation mechanism. This mechanism contributes to pattern formation via the preferential removal of oxygen, resulting in the creation of carbon clusters with a graphite-like composition^29^. Remarkably, after Ar^+^ treatment, the bulk properties of the polymer remain unaffected, including its wetting properties^29^. Compared to other nanopatterning techniques such as photolithography, ion-induced nanopatterning is performed directly on the material and completed within a few minutes, does not require masks, and does not produce toxic byproducts.

When tested against microorganisms with marked differences in cell wall composition and size (e.g., *E. coli, B. subtilis, S. aureus*, and mammalian cells), we found that the nanospikes in the nanostructured BC disrupted the envelope preferentially in bacterial cells, but did not compromise the integrity of murine pre-osteoblasts (MC3T3-E1). Importantly, bacterial envelope rupture was a time-dependent event that was also affected by the geometry and spacing of the nanostructures, and the size and mechanical properties of the microorganisms. For example, the time in contact with the nanotopography dramatically influenced the percentage of viable bacteria, which decreased significantly over time. Expressly, the percentage of viable *E. coli* and *B. subtilis* declined substantially at 24 h despite the accumulation of bacterial debris at the material interface.

This is an important feature, given that the risk of device rejection due to bacterial contamination is highest during the first 24 h following biomaterial implantation^3^. Furthermore, because the nanospikes were cytocompatible towards murine preosteoblasts, we believe the integration of the material into the host tissue can also provide additional protection against infection.

Interestingly, both CoArBC and LgArBC were almost depleted of viable *B. subtilis* at 24 h, while the percentage of live *E. coli* decreased from about 98% to nearly 60% in the CoArBC during the same period. Such time-dependent rupture behavior has also been found experimentally in synthetic lipid membranes^41^, red blood cells^42^ and mammalian cells in contact with silicon nanowires and nanostraws^35^, and has been described in the framework of activation energy theory^37,41^. According to this formulation, the breakup of biological membranes is a dynamic process that depends on the magnitude and duration of the applied force. At low loading forces, for example, a membrane will develop fatal nanopores if stressed for a sufficient length of time ^41,42^. The higher survival of *E. coli* towards the nanostructured BC could also stem from enzymatic degradation of the cellulose. However, we believe this scenario was unlikely because, after Ar^+^ treatment, the BC developed clusters rich on C-C bonds of few nanometers in thickness, which are hard to break even with energetic ions^29^.

We also found that the local geometrical features and spacing of the nanospikes were critical parameters in the mechanical disruption of the bacterial envelope. Specifically, nanospikes with long apexes and low density like those in CoArBC were more effective at killing *B. subtilis* and *E. coli*, compared to denser ones in the LgArBC. Those results are markedly different from those reported by Kelleher and coworkers^43^, who found that tightly packed nanopillars had higher bactericidal activity against *Pseudomonas fluorescens.* However, *S. aureus* was unaffected by the underlying topography and incubation time, probably because this bacterium was able to colonize the spacing separating adjacent nanostructures, particularly on CoArBC. Remarkably, we also found that *B. subtilis* were killed more efficiently than *E. coli* cells. This finding is in sharp contrast with previous reports using analogous nanotopographies, which have suggested that Gram-negative strains like *E. coli* are more susceptible to mechanical rupture than their Gram-positive counterparts^14^. However, our findings are consistent with the view that the bactericidal activity of nanoscale protrusions is independent of the material’s chemistry. Our results also agree with the higher antibacterial activity toward Gram-positive exhibited by other low-dimensional nanomaterials, including single-walled carbon nanotubes and graphene oxide nanowalls ^44,45^. Importantly, based on experimental evidence and computational simulations, those low-dimensional nanomaterials are believed to inflict membrane rupture by mechanical means as a consequence of direct physical contact between the bacterial membrane and the material, and as a result of the increased charge transfer^8,45^.

Collectively, the data support a model whereby the bactericidal activity of the nanostructured celluloses is a function of the cell’s adherence time, nanospike’s geometry and spacing, cell shape, and mechanical properties of the cell wall. Of the possible mechanisms by which bactericidal nanotograhies can inflict bacterial envelope damage, mechanical stretching resulting from a localized pressure appears to bear the most evident link. In a pressure-induced mechanism, one may expect a thick membrane to deform less and, thus, experience higher localized stress at small elements of the envelope, increasing the likelihood of nanostructure penetration. In contrast, softer membranes like those in Gram-negative bacteria and mammalian cells behave as resilient viscoelastic shells, redistributing the applied stress and thus, requiring a higher external force for rupture^36^. Indeed, this viscoelastic behavior has been reported previously for *E. coli*, which could regulate cell wall synthesis upon compression^46,47^, and adapt its growing morphology into irregular shapes when confined in narrow microcavities, while *B. subtilis* was unable to do so^48^. Similarly, mammalian cells have been found to conform and wrap around HAR silicon nanowires, with a probability of nanowire penetration being very low because of the cell’s membrane deformability. Also, we believe that a pressure-induced mechanism may explain why nanospikes with the same aspect ratio, Young’s modulus, and surface chemistry but different spacing have a very distinct bactericidal activity. In particular, a dense nanospike array like that in LgArBC could distribute the applied stress across the cell surface, reducing the material’s bactericidal activity. However, this attribute may also be lost if the spacing is larger than the microorganism size, as in the case of *S. aureus* in contact with CoArBC.

To unveil the mechanism responsible for the preferential loss of integrity in *B. subtilis*, we looked for quantifying the critical loading force to penetrate *E. coli* and *B. subtilis* using cantilever tips with radii close to those in CoArBC and LgArBC. We found that forces below 3 nN were enough to breakthrough *E. coli* and *B. subtilis*, which agrees with the breakthrough forces reported for *Salmonella Typhimurium*^40^. However, *B. subtilis* required less force on average to be penetrated compared to *E. coli.* Most remarkably, these breakthrough forces agree very well with published mathematical predictions about the penetration forces of HAR nanostructures on biological membranes with Young’s modulus in the range of 40-150 MPa^36^, which are within the stiffness estimated for *E. coli* and *B. subtilis* by other authors (23±8 MPa and 49±20 MPa for *E. coli* cells in the axial and circumferential directions respectively^49^, and 100-200 MPa for *B. subtilis*^50^). Indeed, the force required to penetrate a stiff membrane (150 MPa) was computed to be ≈ 0.88 nN, while a soft membrane (40 MPa) needed a more significant penetration force and equal to ≈ 1.42 nN^36^, in excellent agreement with our experimental measurements. The indentation depth measured by AFM was also in quantitative consonance with membrane deformation observed by performing serial cross-section at the bacterium/nanotopography interface, further suggesting that membrane rupture by a localized pressure was highly probable.

Remarkably, both *E. coli* and *B. subtilis* retained its integrity upon repeated indentation. This observation agrees with previous experiments performed on *S. typhimurium*^40^, which recovered after multiple puncturing. However, unlike the scenario where the bacterial envelope can recover upon nanoprobe retraction, we believe the nanospikes can compromise the integrity of bacteria by their capacity to cause permanent cell wall deformation and remain inside the cell. Permanent deformation of biological membranes has been reported in red blood cells using micropipette aspiration by applying to small but sustained forces for an extended period^42^. Unexpectedly, the breakthrough force did not increase significantly by indenting *E. coli* with a cantilever with a larger tip radius (30±9 nm), in consonance with published results of indentation-induced membrane rupture in bacteria and mammalian cells using AFM nanoprobes^38,40^. Recent work has proposed that such an unexpected behavior originates from two types of forces acting on a lipid membrane upon indentation: short-ranged forces that elastically deformed the membrane, and long-range entropic forces that result from the confinement of thermal undulations. While rupture via short-ranged forces is a direct consequence of the applied pressure, thermal undulations depend on the size of the membrane in contact with the nanoprobe tip, and independent of tip geometry^51^.

In summary, we conferred bactericidal properties to BC hydrogel by only altering its topography at the nanoscale and revealed new insights about the biophysical mechanism underlying this phenomenon. Specifically, we obtained two sets of HAR nanostructures with a “nanospike” appearance of approximately the same aspect ratio, but different size and spacing. When tested against microorganisms with marked differences in cell wall composition, we found that the bactericidal activity of the nanospikes in BC was affected by the bacterium’s adherence time, nanospike geometry and spacing, and the size and mechanical properties of the microorganisms, pointing out a pressure-induced mechanism. Most importantly, we found that these nanospikes inflicted more significant membrane damage in *B. subtilis* than in *E. coli*. By mimicking the loading forces to breakthrough *E. coli* and *B. subtilis*, we revealed that forces below 3 nN were enough to penetrate both *E. coli* and *B. subtilis*, but *B. subtilis* required less force, supporting our experimental observations and previous mathematical predictions of membrane rupture by a localized pressure. Even though our findings are in sharp contrast with previous reports on bactericidal nanotopographies, those are in line with the higher bactericidal susceptibility of Gram-positive bacteria observed in other low-dimensional materials, hence breaching the gap between bactericidal nanotopographies and colloidal suspensions. Remarkably, the nanospikes in BC did not compromise the integrity of murine pre-osteoblasts, suggesting the material’s biocompatibility towards mammalian cells, a requisite for tissue restoration. Moreover, this feature may also provide additional protection against harmful bacterial contamination by integrating the material to the host tissue. We believe that our findings will provide a better understanding of the mechanobiology of bacterial cells at the interface with nanoscale structures, which is fundamental for the rational design of bactericidal interfaces.

## 4. Materials and Methods

### Nanostructured bacterial cellulose

Two sets of never-dried bacterial cellulose (BC) were used for nanopatterning. The first set was grown in the laboratory by culturing *Komagataeibacter xylinum* (ATCC® 700178) in a culture medium containing yeast extract, peptone, and mannitol at 30 °C for 7 days as described by Arias and coworkers^52^. This BC was designated as lab-grown bacterial cellulose (LgBC). The second set of BC pellicles were purchased from Jenpolymers (Germany), and which correspond to pellicles produced by *K. xylinum* DSM 14666, and named here as commercially available bacterial cellulose (CoBC). Both sets of never-dried BC were cut in pieces of 0.25 cm^2^ and dried at room temperature on glass slides of the same size. Air-dried BC samples were then inserted in a vacuum chamber and irradiated with argon ions (Ar^+^) at a fluence of 10×10^17^ cm^-2^ using ion energy of 1 keV at normal incidence. The base pressure of the vacuum chamber was 5×10^−6^ Pa, and ion beam irradiation was performed at a gas pressure of 2 × 10^−4^ Pa. The operating current was kept between 0.2 to 0.3 mA.

### Young’s modulus determination of nanostructured bacterial cellulose

The Optics11 Piuma soft material indenter was used to determine the Young modulus of pristine and argon-treated BC. All the materials were indented in aqueous solution (ultrapure water) with a spherical glass bead of 23 µm in diameter and stiffness of 4.02 N/m. Up to 25 indentations were performed per sample, with a separation of 50 μm between indentation points. One-way ANOVA with two-pair two *t-*test was performed to determine significant differences between samples at 0.05 level of significance.

### Strains, culture conditions and live/dead staining

Frozen stocks of *Escherichia coli* ATCC^®^25922 were inoculated in Tryptic soy broth and grown overnight to stationary growth phase (OD_630_≈0.8-1) at 37°C with shaking. Similarly, frozen stocks *Staphylococcus aureus* ATCC^®^12600 and *Bacillus subtilis* ATCC^®^6051 were inoculated in Nutrient broth and grown overnight and for 24 h, respectively, at 37°C with shaking (OD_630_≈0.8). For cell adhesion experiments lasting 1h, bacterial suspensions for each strain were prepared by diluting the starter culture in minimum medium M63 supplemented with 20% glycerol (J910500G, Fisher Scientific) to 1:10 dilutions for *E. coli* and *S. aureus* and 1:2 dilutions for *B. subtilis* (*B. subtilis* ATCC^®^6051 exhibited a slower cell growth than the other two strains). M63 minimal media was used because bacteria have been reported to increase biofilm production under low nutrient or starvation conditions^53,54^. For biofilms experiments lasting 24 h, dilutions of 1:100 were made for *E. coli* and *S. aureus* and 1:10 for *B. subtilis*. Each bacterial dilution was incubated in the presence of the pristine and nanopatterned samples in a 48-well plate at 30°C without shaking for 1 h or 24 h. At the end of the incubation period, experimental samples containing attached bacteria were rinsed with phosphate buffer saline (PBS) three times to remove loosely attached cells and stained with LIVE/DEAD™ *Bac*Light™ Bacterial Viability Kit according to the manufacturer’s instructions (ThermoFisher Scientific). Samples were always kept PBS, even during rinsing and imaging, to prevent cell lysis from capillary forces. Live and dead bacteria were imaged at a magnification of 40X and 63X using oil immersion in a confocal laser scanning microscope (Leica SP8), and cell counts were normalized to the total number of cells per unit area. At least two independent experiments were performed using triplicate samples, and 20 to 30 images were taken per condition at random locations. One-way ANOVA with two-pair two *t-* test was performed to determine significant differences between samples at 0.05 level of significance.

### Biofilm quantification via crystal violet

Four to five samples seeded with bacterial dilutions and cultured for 24 hours at 30 C without shaking as described above, were collected at the end of the incubation period and rinsed several times with PBS to remove non-adherent bacteria. Subsequently, samples were stained with 0.1% crystal violet for 5 min. After that, the colorant was discarded, the samples were rinsed up to seven times with ultrapure water and air-dried for 1 hour at room temperature. Then, the samples were treated with pure ethanol for 5 min to solubilize the biofilms. The solubilized colorant was collected and transferred to a 96 well-plate. The absorbance was read at 570 nm. One-way ANOVA with two-pair two *t-*test was performed to determine significant differences between samples at 0.05 level of significance.

### Scanning electron microscopy and focused ion beam

Pristine and nanostructured BC were seeded with the bacterial suspension at the dilutions indicated previously for 1 h. Subsequently, samples were rinsed with PBS two times to remove weakly attached bacteria and subsequently fixed with 10% formalin solution for 1 h at room temperature. Samples were then washed with ultrapure water and serially dehydrated for 10 min each with ethanol dilutions prepared at 20%, 40%, 60%, and 80%. After that, samples were left on 100% ethanol and placed inside a desiccator connected to a vacuum line. Dried samples were sputter-coated with a thin-film of Au-Pd and imaged at 5 kV with a 10 mM current using a Hitachi 4800 scanning electron microscope. Cross-sections of the same samples were performed in a Scios 2 Focused Ion Beam microscope. To that end, rectangular cross-sections of 25×15 µm in size and 3 µm in depth were made using an ion beam current of 3 nA. Cleaning cross-sections were made using an ion beam current of 0.1 nA. Images were taken using a voltage of 2 kV and probe current of 0.1 nA, dwell time of 10µs, and line integration of 10.

### Cytocompatibility test

MC3T3-E1 Subclone 4 (ATCC^®^ CRL-2593™) preosteoblastic cells were cultured on duplicate samples of tissue-cultured polystyrene (Fluorodish, World Precision Instruments) and commercially available argon-treated BC (CoArBC). Cells were cultured at a density of 5×10^3^ cells/cm^2^ and incubated at 37 C for 24 h using an alpha minimum essential medium (MEM) supplemented with 10% fetal bovine serum. At the end of the incubation period, samples were rinsed with phosphate buffer saline and stained for 15 min with LIVE/DEAD™ Viability/Cytotoxicity Kit (ThermoFisher). Samples were subsequently imaged using a confocal laser scanning microscope (Leica SP8) at a magnification of 40X and 63X using oil immersion. Up to 30 images were taken per condition at random locations.

### Force volume indentation and analysis

Four Pyrex-Nitride Probe Triangular (PNP-TR, NanoWorld) and two NPG Triangular (Bruker AFM Probes) cantilevers with nominal resonance frequencies of 17 kHz and 23 kHz respectively, were used for the indentation experiments. The spring constant and inverse optical lever sensitivity for each cantilever was determined by the thermal vibration spectrum method and was 22.11 ± 0.50 pN/nm and 68.11 ± 5.99 nm/V for the PNP-TR, and 125.40 ± 1.92 pN/nm and 52.45 ± 4.53 nm/V for NPG (B). The characterization of the nanomechanical properties of *E. coli* and *B. subtilis* was performed through a force-volume analysis using an Asylum Cypher Atomic Force Microscope (AFM) and recorded in phosphate buffer saline (PBS) at room temperature. Before the experiment, *E. coli* and *B. subtilis* were grown as previously described, collected by centrifugation at 1500 rpm for 10 min from stationary phase cultures (OD_560nm_=0.8) and resuspended in 200 uL of PBS. A small drop of bacterial suspension was then placed on a clean silicon substrate (squares of 10 mm side), incubated for 30 min, rinsed with PBS three times, and then mounted in a magnetic holder and transferred to the AFM instrument. Immediately after, a small drop of PBS was added to the cantilever and silicon substrate. The experiment was performed in contact mode, at a scan rate of 0.99 Hz and a velocity of 1.98 µm/s. For each cell type, more than 50 force volume maps containing 196 pixels each were generated on random areas of 5×5 µm^2^, in which each pixel corresponded to a force curve acquired with a trigger force of 2 to 5 nN at 1 µm of distance from the surface. The breakthrough events and determination of indentation depth were analyzed on a MATLAB-based application (Force Review Automation Environment or FRAME)^55^. Formalin-fixed *E. coli* seeded on wrinkled polydimethylsiloxane (PDMS) was imaged with a PNP-TR cantilever in contact mode in PBS using a force of 20 pN, and subsequently indented with a trigger force in the range of 2 to 5 nN following a procedure as described previously. Wrinkled PDMS was obtained as described by Arias and coworkers^56^. Determination of the nanoprobes’ radius was performed using a calibration grid (Silicon grating TGT01, NT-MDT Spectrum Instruments) (**Supplementary Figures S5** and **S6**).

### Image and data analysis

We analyzed confocal laser scanning micrographs and scanning electron images using ImageJ. To obtain the percentages of the viable to dead cells, we adjusted the threshold for each image and then used the particle analyzer tool to compute the number of viable and dead cells. For the scanning electron micrographs, the scale bar was set up for each image using a known distance, and subsequently, the size of the nanostructures and other structures were obtained manually using the imageJ’s measure tool.

## Acknowledgments

The mammalian cell viability test was performed with the assistance of Andrea Mesa Restrepo. Material characterization was carried out in the Materials Research Laboratory Central Research Facilities at the University of Illinois at Urbana-Champaign. The laser scanning confocal fluorescence micrographs were acquired at the Microscope Suite of the Beckman Institute for Advanced Science and Technology at the University of Illinois at Urbana-Champaign.

## Contributions

S.L.A conceived the idea. Experiments were designed by S.L.A and J. S. and carried out by S.L.A, J.S., and J.D. The manuscript was written by S.L.A. A.C helped with the analysis and interpretation of results. The principal investigator is J.P.A. All authors edited the manuscript.

## Materials & Correspondence

Correspondence and material requests should be addressed to Sandra L. Arias

