## Supplementary materials for "Bacterial envelope damage inflicted by bioinspired nanospikes grown in a hydrogel"

**This PDF file includes:**

Supplementary Figures **S1** to **S6**

Supplementary References, *57*

**Supplementary Figures**

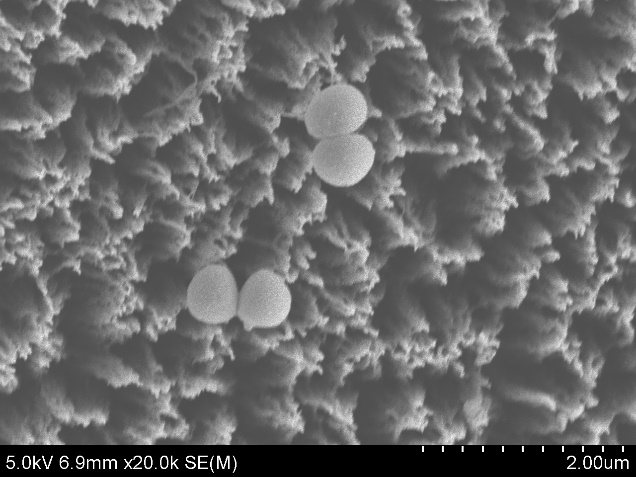

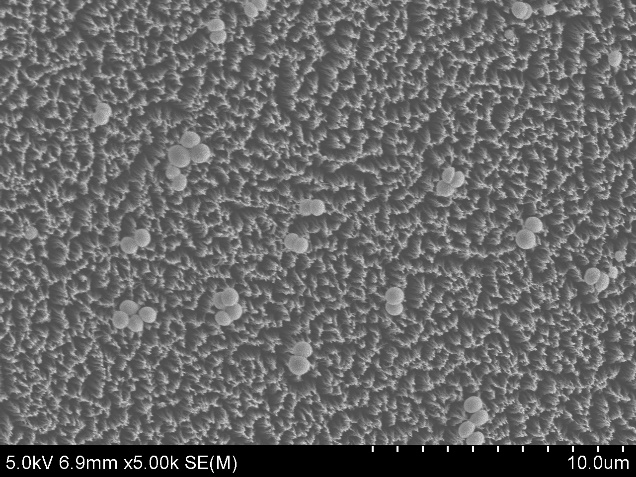

10 µm

2 µm

**a.**

**b.**

**Supplementary Figure S1. (a)** SEM of *S. aureus* cultured on CoArBC. **(b)** High-resolution image of the region indicated by the box in **(a)**. Notice that *S. aureus* could fit inside the spacing separating adjacent nanospikes (red arrows).

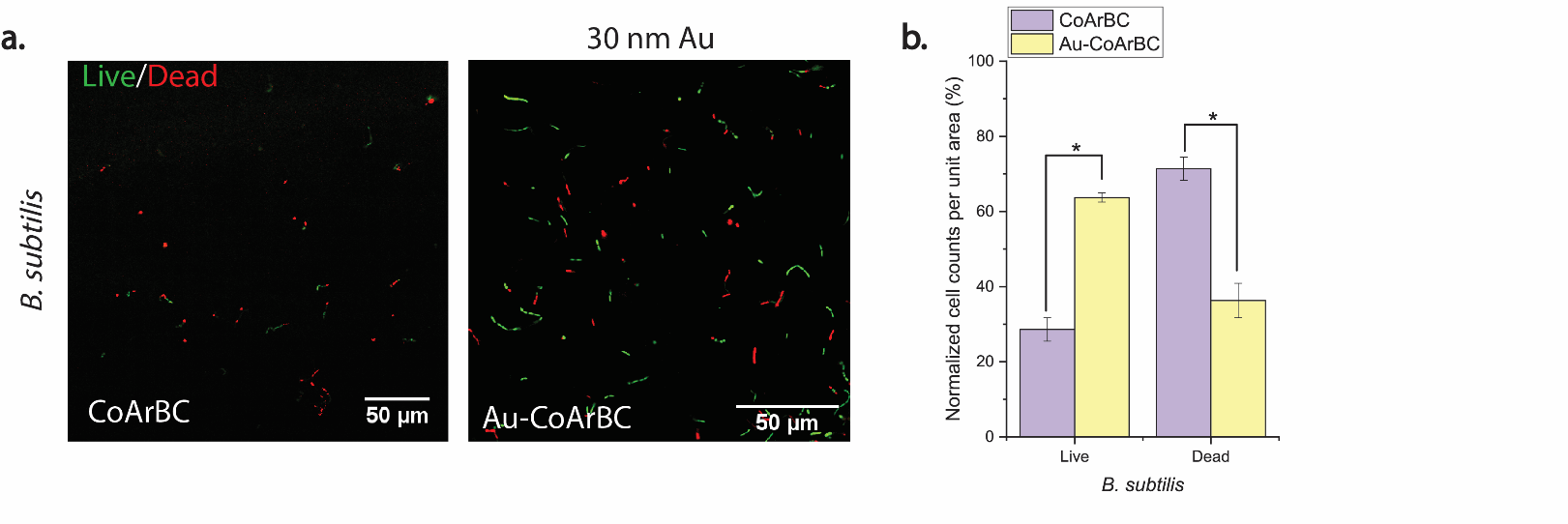

**Supplementary Figure S2. (a)** Live/dead staining of *B. subtilis* seeded on CoArBC and gold-coated CoArBC (Au-CoArBC)**. (b)** Percentage of the live to dead cells normalized to the total number of cells per unit area.

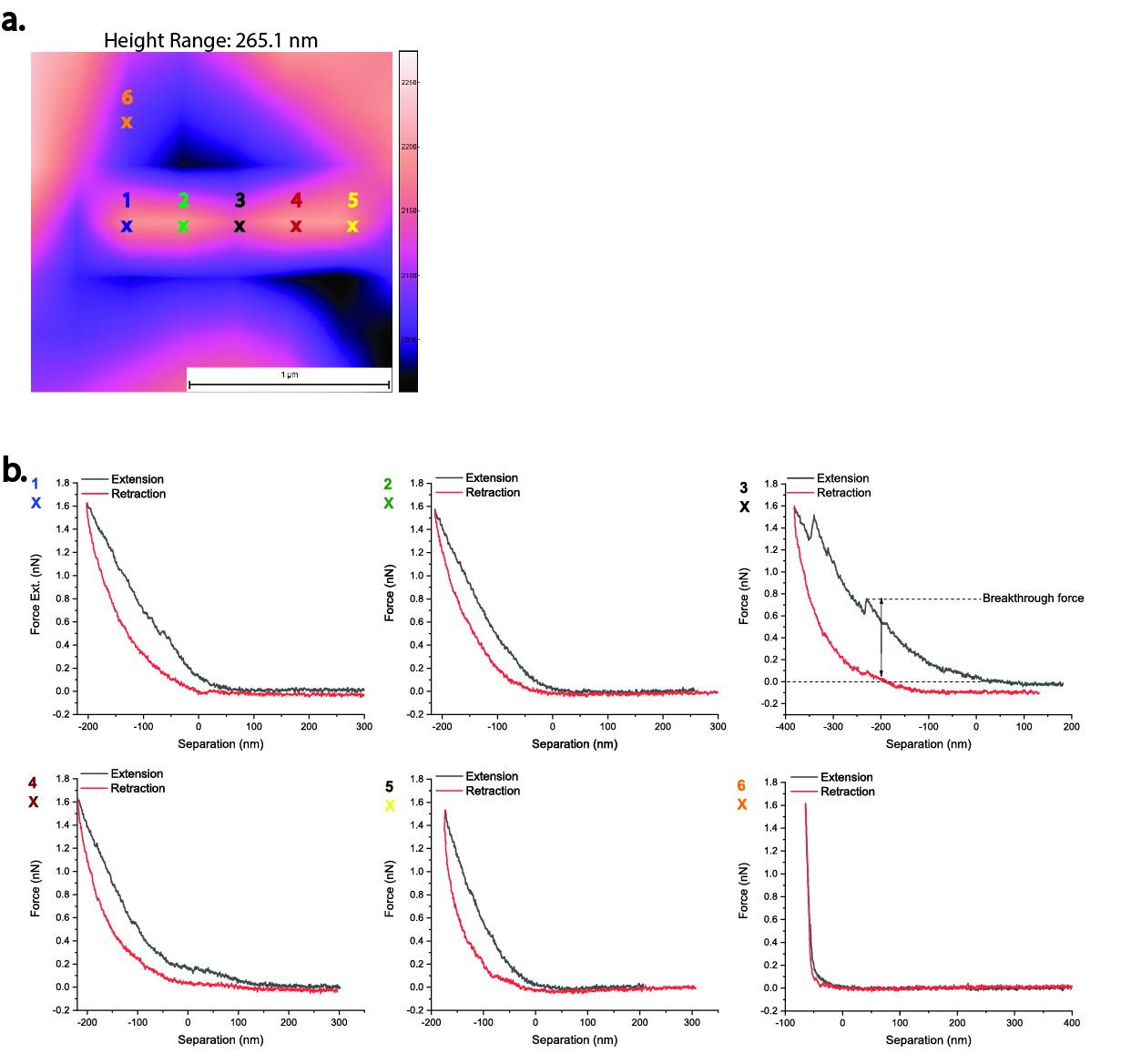

**Supplementary Figure S3. (a)** Single *E. coli* cell immobilized inside wrinkled polydimethylsiloxane (PDMS) and reconstructed using SPIP software from a force volume map image. This cell was indented at five positions along its longitudinal axis, as depicted by the cross marks. Each of these cross marks correspond to a force curve, which are given in **(b).** The force curve for the wrinkled PDMS substrate (cross mark number 6) is provided for comparison purposes. Notice the spatially dependent elasticity of *E. coli* for a loading force of 2 nN.

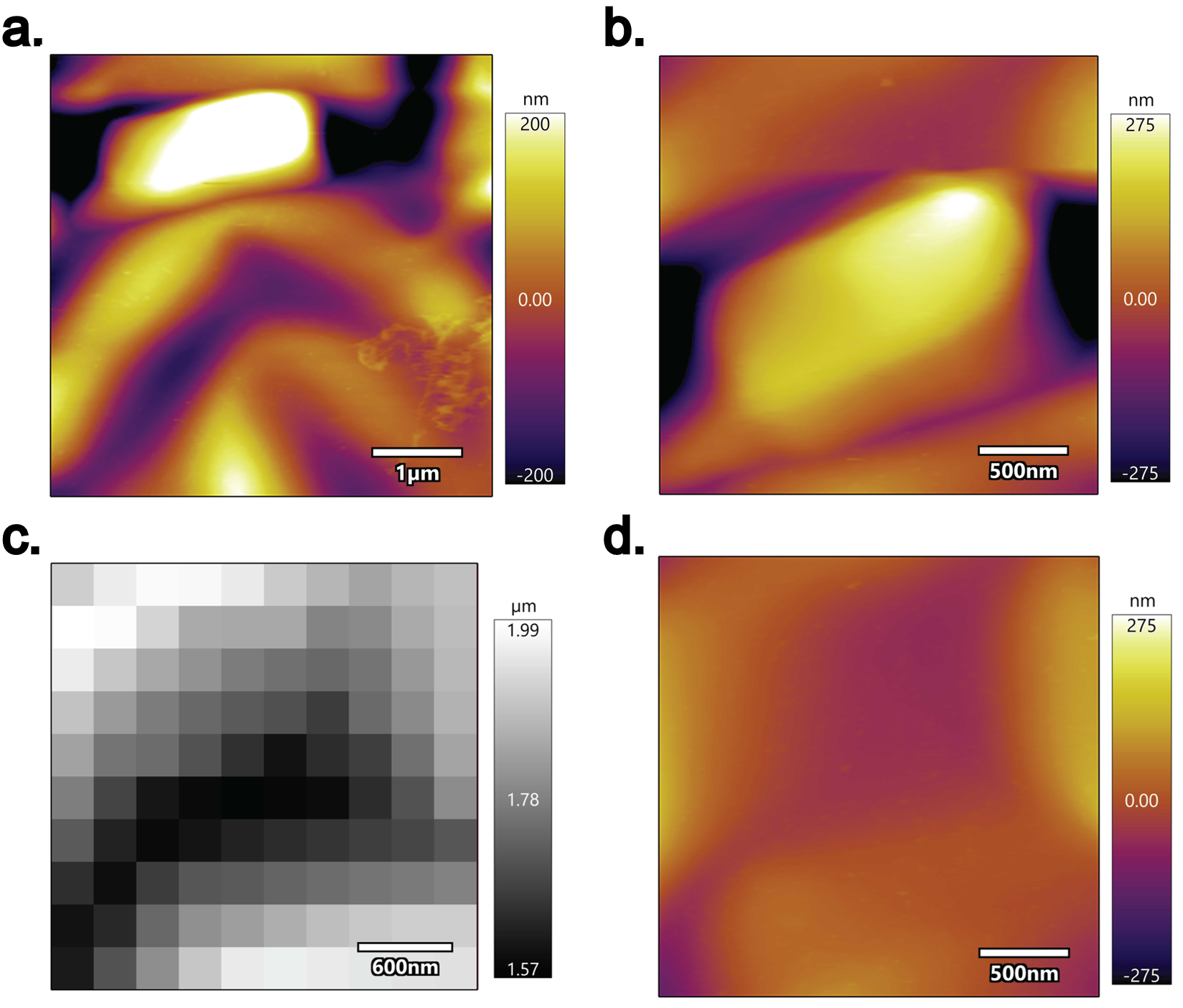

**Supplementary Figure S4. (a)** Fixed *E. coli* cell immobilized inside the wrinkles of a PDMS and image by contact mode AFM. **(b)** High-resolution scan of **(a). (c)** Force volume mapping of **(b),** performed with a loading force of 2 nN. The map shows the depression of the cell as it is indented by a nanoprobe. **(c)** The cell is peeled away from the substrate.

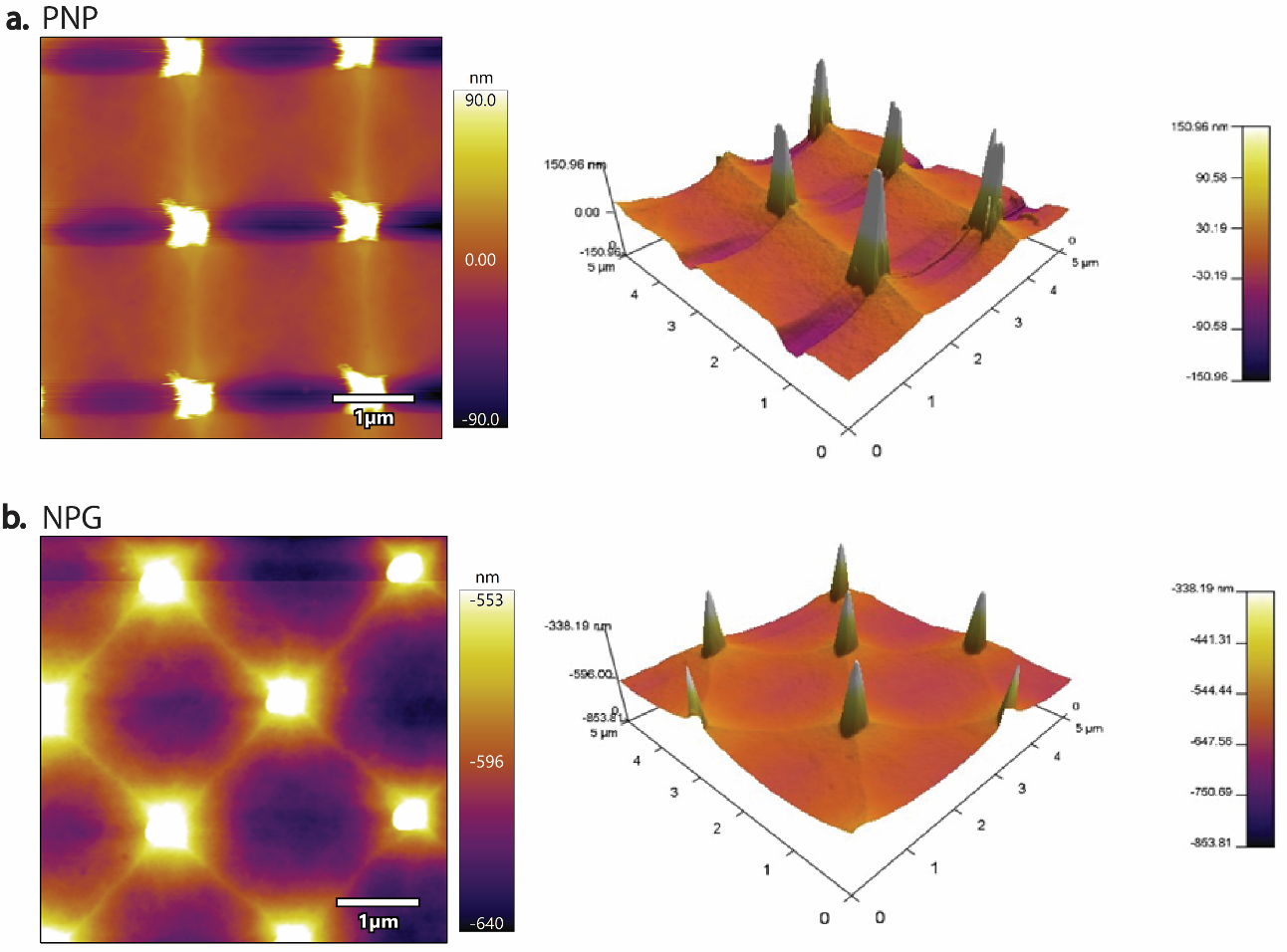

**Supplementary Figure S5.** Representative atomic force microscope (AFM) images of a silicon calibration grid (TGT01, NT-MDT Spectrum Instruments) scanned with a pyrex-nitride probe (PNP) **(a)** and an NPG probe **(b)**. The two-dimensional images on the left were obtained in contact mode using an Asylum Cypher AFM instrument. The panels on the right correspond to a three-dimensional view of the images on the left. The silicon calibration grid has an active area of 2x2mm, tips radii of less than 10 nm, diagonal pitch of 3 µm, square pitch of 2.12 µm, and an angle at the top of the tip of less than 20°.

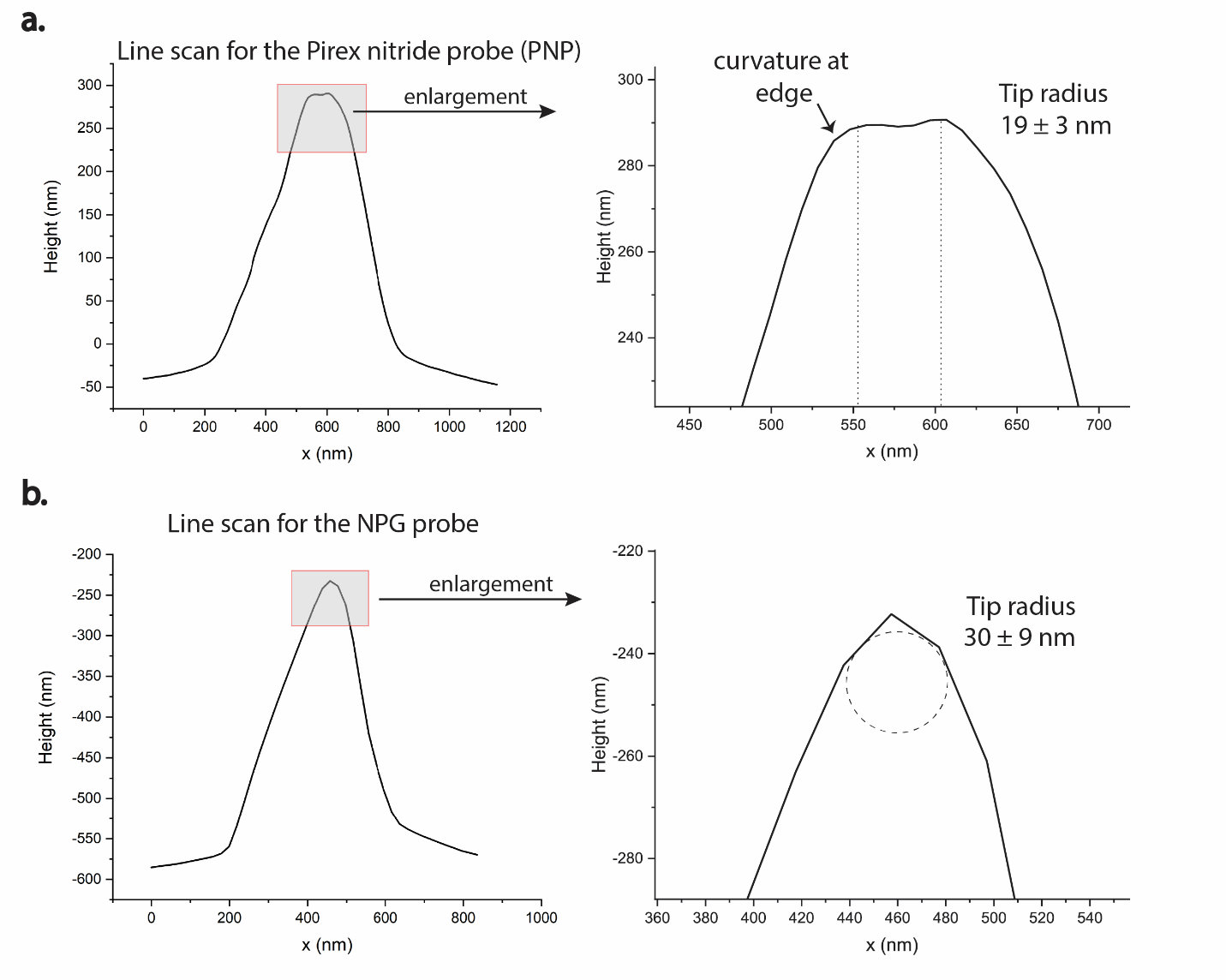

**Supplementary Figure S6.** Determination of a nanoprobe tip radius by extracting the line profiles of calibration grid scans (see **Figure** **S5).** Tip radii were estimated by following the method described by Hübner and coworkers^57^, which states that for nanostructures with large sidewalls and high aspect ratios, the measured sidewalls and edges reflect the shape and tip radius respectively. Here, the estimated tip radii of about 19 and 30 nm for the PNP and NPG, respectively, agree with the nominal values reported by the manufacturers (10 nm for PNP and 30 nm for NPG).
